# Phosphoproteomic Landscape of AML Cells Treated with the ATP-Competitive CK2 Inhibitor CX-4945

**DOI:** 10.1101/2021.01.04.425248

**Authors:** Mauro Rosales, Arielis Rodríguez-Ulloa, Vladimir Besada, Ailyn C. Ramón, George V. Pérez, Yassel Ramos, Osmany Guirola, Luis J. González, Katharina Zettl, Jacek R. Wiśniewski, Yasser Perera, Silvio E. Perea

## Abstract

Casein kinase 2 (CK2) regulates a plethora of proteins with pivotal roles in solid and hematological neoplasia. Particularly, in acute myeloid leukemia (AML) CK2 has been pointed as an attractive therapeutic target and prognostic marker. Here, we explored the impact of CK2 inhibition over the phosphoproteome of two cell lines representing major AML subtypes. Quantitative phosphoproteomic analysis was conducted to evaluate changes in phosphorylation levels after incubation with the ATP-competitive CK2 inhibitor CX-4945. Functional enrichment, network analysis, and database mining were performed to identify biological processes, signaling pathways, and CK2 substrates that are responsive to CX-4945. A total of 273 and 1310 phosphopeptides were found differentially modulated in HL-60 and OCI-AML3 cells, respectively. Despite regulated phosphopeptides belong to proteins involved in multiple biological processes and signaling pathways, most of these perturbations can be explain by direct CK2 inhibition rather than off-target effects. Furthermore, CK2 substrates regulated by CX-4945 are mainly related to mRNA processing, translation, DNA repair, and cell cycle. Overall, we evidenced that CK2 inhibitor CX-4945 impinge on mediators of signaling pathways and biological processes essential for primary AML cells survival and chemosensitivity, reinforcing the rationale behind the pharmacologic blockade of protein kinase CK2 for AML targeted therapy.

## 1. Introduction

Protein phosphorylation is an essential post-translational modification in most cellular processes, making of protein kinases promising therapeutic targets for a wide variety of disorders, including cancer [1, 2]. Among the protein kinases involved in cell signaling networks, casein kinase 2 (CK2) is responsible of about 25% of all cell phosphoproteome [3]. CK2 is a constitutively active and ubiquitously expressed Ser/Thr-protein kinase composed of two catalytic subunits (α or its isoform α’) and two regulatory subunits (β) [4]. The CK2 consensus sequence (pS/pT-x1-x2-E/D/pS/pT, in which x1 ≠ P), is a small motif characterized by several acidic residues in the proximity of the phosphorylatable amino acid, as well as the absence of basic residues in those positions [5]. Concerning CK2 substrates, about one third are involved in gene expression and protein synthesis, while numerous are signaling proteins implicated in cell growth, proliferation, and survival [3, 6]. Also, a small number of CK2 substrates are classical metabolic enzymes or associated with some virus life cycle [3].

Protein kinase CK2 has been linked to basically all the hallmarks of malignant diseases [7, 8]. Accordingly, several CK2 inhibitors have been described, including small organic compounds designed to target the ATP-binding site on the CK2 catalytic subunit, flavonoids and a synthetic cell-permeable peptide termed CIGB-300, originally designed to block CK2-mediated phosphorylation through binding to phosphoacceptor domain in the substrates [9-11]. Additionally, a cyclic peptide that antagonizes the interaction between the CK2 α and β subunits and antisense oligonucleotides that reduce CK2 alpha subunit transcription have also been explored [12, 13]. However, only the ATP-competitive inhibitor CX-4945 and the synthetic peptide CIGB-300 have advanced to human clinical trials in and shall provide proof-of-concept for CK2 as a suitable oncology target [14, 15].

Acute myeloid leukemia (AML) is one of the most frequent hematologic malignancies and high-expression of CK2α subunit has been connected to a worse prognosis in AML patients with normal karyotype [16, 17]. Actually, CK2 is implicated in multiple signaling pathways, all of them essential for hematopoietic cell survival and function, and leukemic cells have been demonstrated to be more sensitive to downregulation of protein kinase CK2 [18, 19]. The latter becomes particularly relevant since AML stand among the most aggressive and lethal types of cancer and are often characterized by resistance to standard chemotherapy as well as poor long-term outcomes [20].

In recent years, quantitative phosphoproteomic approaches have been useful to explore the cellular response to kinase inhibition in different types of cancer cells [21]. In fact, the proteomic and phosphoproteomic patterns associated with prognosis of AML patients and its progression from diagnosis to chemoresistant relapse has been recently described, studies that suggested the importance of CK2 for chemosensitivity in human AML primary cells [22, 23]. Besides, the CK2-dependant phosphoproteome has been explored by quantitative phosphoproteomic using not only CK2 inhibitors in HEK-293T, HeLa, and NCI-H125 cells, but also through genetic manipulation of CK2 subunits in C2C12 cells [24-27]. However, the impact of CK2 inhibition has not been widely assessed in AML cells, since to our knowledge no previous phosphoproteomic studies have been conducted for CK2 inhibitors in this particular hematological pathology. Considering the above, we decided to explore the CK2-regulated phosphoproteome and the consequent signaling networks perturbations induced after exposure of AML cells to CK2 inhibitor CX-4945. Mass spectrometry (MS)-based phosphoproteomics profiling allowed us to gauge the global impact of CX-4945 in human cell lines representing two differentiation stages and major AML subtypes.

## 2. Materials and Methods

### 2.1. Cell Culture and AlamarBlue Assay

Human AML cell lines HL-60 and OCI-AML3 were originally obtained from the American Type Culture Collection (ATCC, VA, USA) and the German Collection of Microorganisms and Cell Cultures (DSMZ, Braunschweig, Germany), respectively. Both cell lines were cultured in RPMI 1640 medium (Invitrogen, CA, USA) supplemented with 10% (v/v) fetal bovine serum (FBS, Invitrogen, CA, USA) and 50 µg/mL gentamicin (Sigma, MO, USA) under standard cell culture conditions. The antiproliferative effect of CX-4945 on HL-60 and OCI-AML3 was assessed using AlamarBlue assay (Life Technologies, CA, USA). Briefly, cells were seeded in flat-bottom 96-well plates (2 × 10^5^ cells/mL, 200 µL/well) and 24 h later serial dilutions 1:2 ranging from 50-1.6 µM of CX-4945 (Selleck Chemicals, TX, USA) were added. After 72 h of incubation, AlamarBlue was added at 10% (v/v), and cell suspension were incubated for 4 h. Next, fluorescence was measured using a CLARIOstar microplate reader (BMG LABTECH, Ortenberg, Germany) and half-inhibitory concentration (IC_50_) values were estimated using CalcuSyn software (v2.1) (Biosoft, Cambridge, United Kingdom).

### 2.2. Sample Preparation and Phosphopeptide Enrichment

HL-60 and OCI-AML3 cells (10^7^ cells per each condition, three biological replicates) were treated or not with 5 µM CX-4945 (Selleck Chemicals, TX, USA) for 8 h. After collected by centrifugation and washed with PBS, cells were resuspended in lysis buffer containing 2% SDS and 50 mM DTT. Samples were boiled at 95 °C for 10 min and proteins were extracted by multienzyme digestion filter-aided sample preparation (MED-FASP) with overnight lys-C and tryptic digestions [28]. Phosphopeptides were then enriched from each digestions using TiO_2_ beads as previously described [29]. For enrichment, ‘Titansphere TiO_2_ 10 µm’ (GL Sciences, Inc., Tokyo, Japan) was suspended in 200 µL of 3% (m/v) dihydroxybenzoic acid in 80% (v/v) CH_3_ CN, 0.1% CF_3_ COOH and diluted 1:4 with water and later used at a 4:1 ratio (mg beads: mg peptides). Next, 2 mg TiO_2_ (per mg peptides) was added to each sample and incubated at room temperature under continuous agitation for 20 min. The titanium beads were sedimented by centrifugation and the supernatants were collected and mixed with another portion of the beads and incubated as above. The bead-pellets were resuspended in 150 µL of 30% (v/v) CH_3_ CN containing 3% (v/v) CF_3_ COOH and transferred to a 200 µL pipet tip plugged with one layer of Whatman glass microfiber filter GFA (Sigma, MO, USA). The beads were washed 3 times with 30% (v/v) CH_3_ CN, 3% CF_3_ COOH (v/v) solution and 3 times with 80% CH_3_ CN (v/v), 0.3% CF_3_ COOH (v/v) solution. Finally, the peptides were eluted from the beads with 100 µL of 40% CH_3_ CN (v/v) containing 15% NH_4_ OH (m/v) and were vacuum-concentrated to ∼4 µL. Phosphopeptides were further desalted by Stage procedure [30].

### 2.4. NanoLC-MS/MS and Data Analysis

Chromatographic runs for phosphopeptides and non-phosphopeptides were in home-made column (75 mm ID, 20 cm length). For phosphopeptides, was used a gradient from 5% buffer B (0.1% formic acid in acetonitrile) up to 30% in 45 min, then increase to 60% in 5 min, and up to 95% in 5 min more. Meanwhile for non-phosphopeptides the gradient started at 5% buffer B up to 30% in 95 min, then increase to 60% in 5 min, and up to 95% in 5 min more. An EASY-nLC 1200 system coupled to a QExactive HF mass spectrometer (both from Thermo Fisher Scientific, MA, USA) was used with the nanocolumn being at 60 °C. Peptides were detected in the mass range 300-1650 m/z using data-dependent acquisition and each mass spectrum was obtained at 60000 resolution (20 ms injection time) and followed by 15 MS/MS spectra (28 ms injection time) at 15000 resolution. Identification of peptides and proteins was based on the match-between-runs procedure using MaxQuant software (v1.6.2.10) [31], and considering oxidation (M), deamidation (NQ), N-terminal acetylation (proteins) and phosphorylation (STY) as variable modifications. None fixed modifications were considered as cysteines were not modified. Alignment of chromatographic runs were allowed with default parameters (20 min time window and a matching of 0.7 mins between runs). Filtering and quantification of phosphopeptides were performed in Perseus computational platform (v1.6.2.2) [32]. Reverse and potential contaminant hits were removed, while only phosphosites with localization probability above 0.75 were retained for further analysis. Student’s t Test was employed to identify statistically significant changes (*p*-values lower than 0.05) in phosphorylation and protein levels, after filtering for two valid values in at least one group. An additional fold-change (treated vs. control) cutoff of 1.5 was also applied.

### 2.6. Enrichment Analysis and Sequence Logo

Biological processes significantly represented in differentially-phosphorylated proteins were identified through functional annotation and enrichment analysis, based on the information annotated in the Gene Ontology (GO) database (http://www.geneontology.org/) [33, 34]. Analysis was performed with DAVID (v6.8) web-based tool (http://david.ncifcrf.gov/) and all identified phosphoproteins dataset was used as background [35, 36]. DAVD computes EASE-score, a modified Fisher Exact Test to identify significant enriched biological processes (*p*-values lower than 0.1) [35, 36]. The resulting list of GO terms with its corresponding *p*-values was further submitted to REViGO (http://revigo.irb.hr/) for redundancy reduction [37]. In addition, sequence logos for down-regulated phosphopeptides were generated using WebLogo (v3.6.0) (http://weblogo.threeplusone.com/) and MaxQuant amino acid sequence window was used as input [38].

### 2.7. Enzyme-Substrate Relationship and Kinome Network Analysis

Enzyme-substrate-site relations were retrieved using the integrated protein post-translational modification network resource iPTMnet [39]. iPTMnet is based on a set of curated databases like PhosphoSitePlus (http://www.phosphosite.org) and PhosphoEML http://phospho.elm.eu.org), which annotate experimentally observed post-translational modification [40, 41]. Besides, the KEA2 web tool (https://www.maayanlab.net/KEA2/) was used, first to retrieve information about kinases responsible for phosphoproteome modulation after CK2 inhibition, and second to identify which of such kinases were enriched based on the phosphoproteomic profile [42]. KEA2 is based on an integrative database of kinase-substrate interactions derived from disparate source including literature [42]. The software computes a Fisher Exact Test to distinguish significant enriched kinases (*p*-values lower than 0.05), through statistical analysis [42]. To represent the kinome network, the interactions among the protein kinases associated to the phosphoproteomic profile, according to KEA2 and iPTMnet annotations, were retrieved using the Metascape gene annotation and analysis resource (http://metscape.org) [43]. Such bioinformatics software compiles the information from different integrative databases and applies the MCODE algorithm to extract highly connected regions or complexes embedded in proteins networks [44].

### 2.8. Identification and Analysis of CK2 Substrates

In addition to *bona fide* CK2 substrates, we searched for candidate substrates based on: 1) the presence of the CK2 consensus sequence (pS/pT-x1-x2-E/D/pS/pT, x1 ≠ P) [5], 2) the enzyme-substrate predictions retrieved from NetworKIN database [45], 3) the dataset of high confidence CK2 substrates reported by Bian *et al*. [46] and 4) the phosphoproteins which interact with CK2 according to Metascape database information [43]. Substrates that met at least two of such criteria were selected as the most reliable for further functional analysis. All identified substrates (*bona fide* and putative) were represented in a network context and classified according to biological processes annotated in GO database [33, 34], and the STRING database (http://string-db.org/) was used to identify interactions between proteins [47]. In such analysis only databases and experimental evidences were used as source of interaction data and the confidence score was fixed at 0.4. All protein-protein interaction networks (kinome network and CK2 substrates network) were visualized using Cytoscape software (v.3.5.0) [48].

## 3. Results and Discussion

### 2.1. Profiling the CX-4945-Responsive Phosphoproteome in AML Cells

Advances in high throughput technologies and bioinformatic tools for subsequent data analysis, make possible to explore on a wide-scale fashion the cellular response to inhibition of protein kinases. Particularly, phosphoproteomic studies provide solid evidences regarding kinase-substrates and kinases-kinases relationships involved in the complexity of networks regulating cellular processes in health and disease. Hence, we decided to explore the CK2-regulated phosphoproteome in AML cells using MS-based phosphoproteomic analysis of HL-60 and OCI-AML3 cells treated or not with 5 µM of the CK2 inhibitor CX-4945 during 8 h (**Figure 1A**). Of note, the inhibitory effect of CX-4945 over CK2 enzymatic activity has been previously evidenced by reduction of *bona fide* CK2 substrates phosphorylation and immunoblotting with antibody against pan-CK2 phosphorylated motif [25, 49]. In addition, as measured using AlamarBlue assay, CX-4945 showed a similar dose-dependent inhibitory effect on HL-60 and OCI-AML3 cells proliferation, with IC_50_ values of 7.49 ± 1.55 µM and 4.69 ± 1.59 µM, respectively (**Figure S1**). AML is a highly heterogenous disease, and selected cell lines derive from the most common AMLs (i.e. acute promyelocytic and acute myelomonocytic leukemia), together accounting for roughly two thirds of all AML cases [50]. Moreover, in spite of the similar antiproliferative effect exerted by CX-4945 in both AML cell lines, HL-60 has been described as refractory to CX-4945-induced apoptosis [51]. Thus, selected cells lines not only represent major AML subtypes, but also different niches that can be found in the clinical setting considering its differential sensitivity to CK2 inhibition with CX-4945.

**Figure 1.**
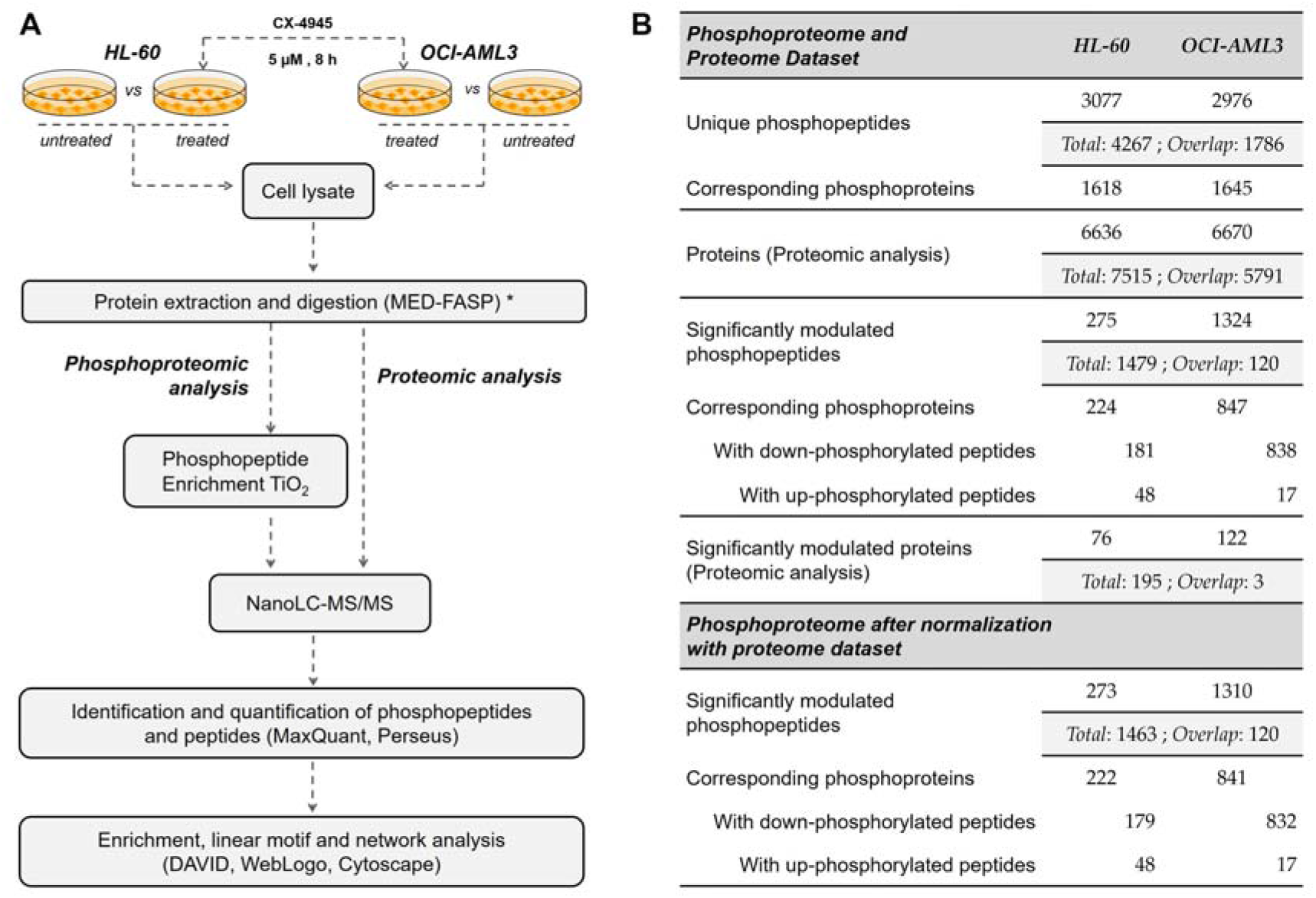
Phosphoproteomic and proteomic analysis of human AML cells treated with the CK2 inhibitor CX-4945: (**a**) Workflow for the exploration of phosphorylation changes induced in HL-60 and OCI-AML3 cells after treatment with CX-4945. Three biological replicates of each group were evaluated; (**b**) Number of identified and significantly modulated phosphopeptides and proteins in each AML cell line. Phosphoproteomic results are showed before and after normalization with the proteome dataset. (*) MED-FASP: multienzyme digestion filter-aided sample preparation [28].

Using this experimental approach, phosphoproteomic analysis of HL-60 led to identification of 3365 phosphopeptides corresponding to 3077 unique phosphopeptides (90% pS, 9.8% pT and 0.2% pY) on 1618 phosphoproteins (**Figure 1B**). Similarly, in OCI-AML3 cells 3177 phosphopeptides were identified, corresponding to 2976 unique phosphopeptides (87.8% pS, 11.9% pT and 0.3% pY) on 1645 phosphoproteins (**Figure 1B**). In parallel, proteomic analysis led to identification of 6636 and 6670 proteins in HL-60 and OCI-AML3, respectively (**Figure 1B**). On the whole, we identified a total of 4267 unique phosphopeptides and 7515 proteins, with 1786 phosphopeptides and 5791 proteins that overlapped between both AML cell lines (**Figure 1B**).

Changes in phosphorylation and protein levels between untreated and CX-4945-treated cells were assessed using Student’s t Test and *p*-value < 0.05 was considered statistically significant. We also applied a fold-change (treated vs. control) threshold of 1.5 (|FC| ≥ 1.5) to define the down- and up-regulated phosphopeptides and proteins. In HL-60 cells 275 phosphopeptides on 224 proteins were significantly modulated, while in OCI-AML3 cells the number was almost 5-fold higher with 1324 on 847 proteins (**Figure 2A, Table S1**). In both cellular contexts, treatment with CX-4945 elicited a global decrease of protein phosphorylation, based on the distribution of down- and up-regulated phosphopeptides in Volcano plots (**Figure 2A**). On the contrary, proteomic analysis indicated that in both cell lines CK2 inhibition showed no bias towards the protein down-regulation (**Figure 2B, Table S2**). Actually, proteome analysis evidenced that changes in phosphorylation upon CX-4945 treatment were mostly independent of protein abundance, since only eight down-regulated proteins (two in HL-60 cells and six in OCI-AML3 cells) had phosphorylation sites significantly inhibited (**Figure 2B**). Those proteins were not considered as differentially phosphorylated after CK2 inhibition, and consequently, were not included in the functional interpretation of the phosphoproteomic profiles.

**Figure 2.**
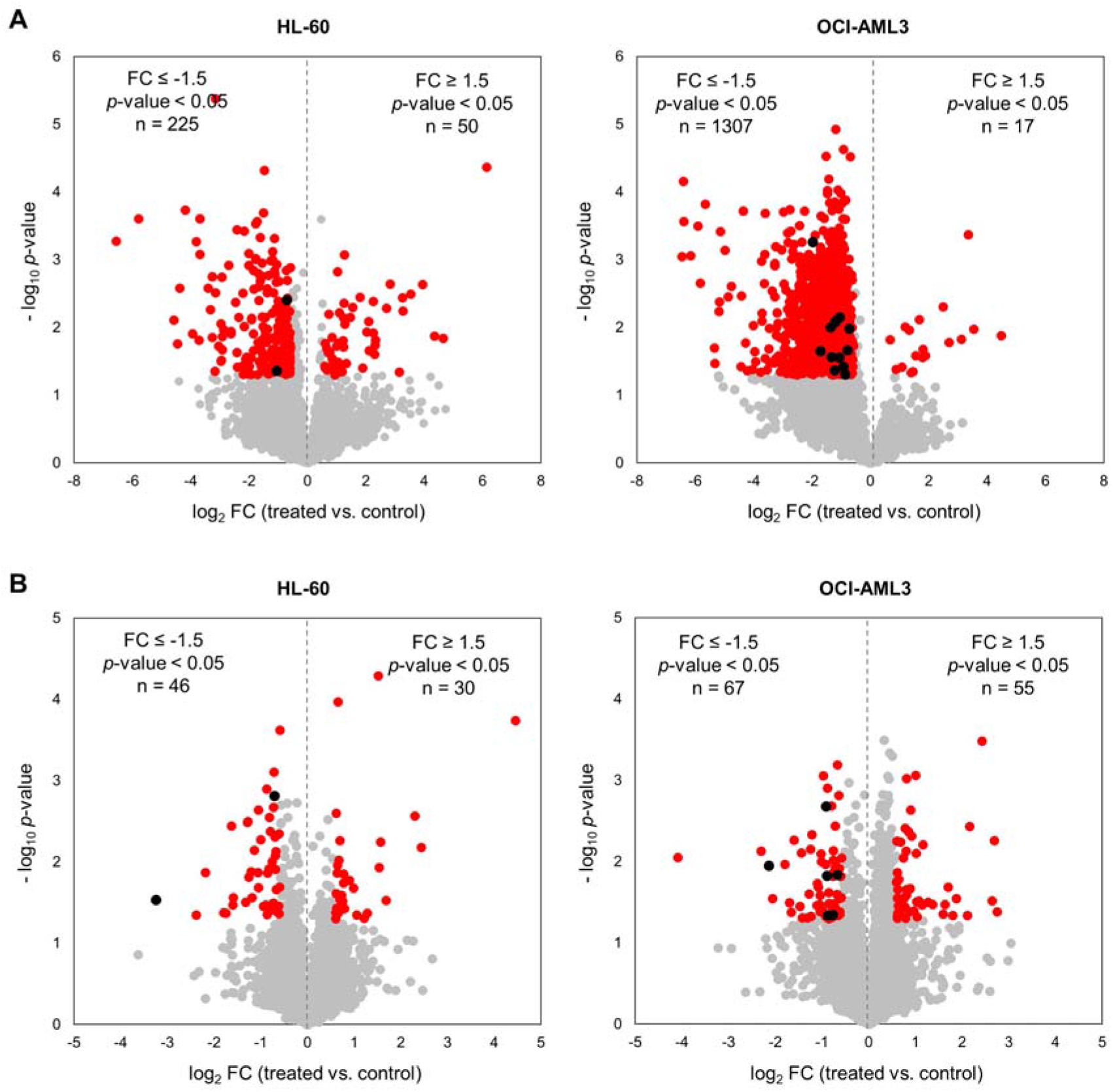
Phosphoproteomic and proteomic profile of human AML cells treated with the CK2 inhibitor CX-4945. Volcano plots of quantified (**a**) phosphopeptides and (**b**) proteins from HL-60 and OCI-AML3 cells after treatment with 5 µM CX-4945 during 8 h. Red points indicate those phosphopeptides/proteins that met statistical significance cut-off (|FC| ≥ 1.5, *p*-value < 0.05). Additionally, black points indicate those phosphopeptides with decreased phosphorylation due to the reduction of the corresponding protein abundance in proteomic analysis (down-regulated proteins are also indicated in black).

In summary, after normalization with the proteome dataset a total of 273 and 1310 significantly modulated phosphopeptides were identified in HL-60 and OCI-AML3 cells, respectively (**Figure 1B, Figure 2A**). Remarkably, such difference indicates that CX-4945 has a more pronounced effect over the CK2-dependant signaling in OCI-AML3 cells, which suggests that the molecular perturbations induced by this inhibitor could rely on the AML cellular background. However, CX-4945 had a similar dose-dependent inhibitory effect on HL-60 and OCI-AML3 cells proliferation (**Figure S1**). It suggests that despite the divergence concerning the molecular impact of CK2 inhibition, there is no differential sensitivity of AML cells towards the overall antiproliferative effect of CX-4945.

### 2.2. Enrichment Analysis of Differentially Modulated Phosphoproteins

For better understanding of putative biological processes perturbed after CK2 inhibition in AML cells, the differentially modulated phosphoproteins were classified in terms of their biological functions using the information from the GO database [33, 34]. Analysis was performed using DAVID web-based tool and GO terms list was further submitted to REViGO for redundancy reduction [35-37]. Significantly represented biological processes in both phosphoproteomics profiles include mRNA processing, regulation of viral process and protein sumoylation (**Figure 3**). Also, phosphorylation sites differentially modulated in HL-60 are located on phosphoproteins related to mRNA splicing, cellular response to DNA damage and ribosome biogenesis, while in OCI-AML3 covalent chromatin modification, nuclear transport, regulation of cell proliferation and gene expression are significantly represented (**Figure 3**). Of note, apoptotic signaling pathway was only identified as significantly enriched in OCI-AML3 cells. Consistently, previous studies have evidenced that HL-60 cell line displays refractoriness to CX-4945 induced apoptosis, probably owing to the absence of p53 protein (HL-60 cells are p53 null) and the lower CK2 protein level and activity in comparison to other AML cell lines [51]. In such study it was demonstrated that CK2 inhibition not only triggers apoptotic cell death in AML cell lines, but also in freshly isolated blasts from AML patients [51].

**Figure 3.**
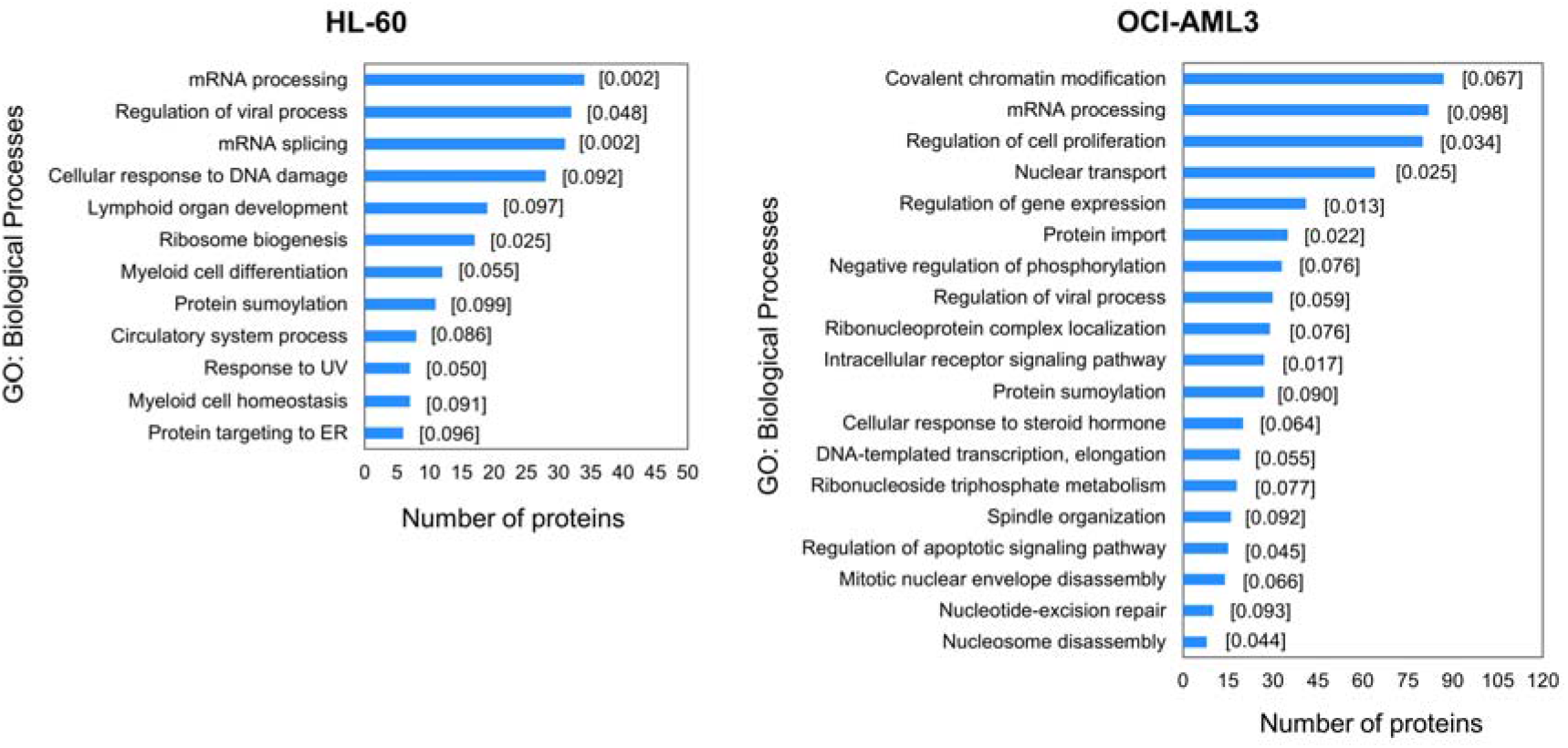
Enrichment analysis for differentially modulated phosphoproteins in HL-60 and OCI-AML3 cells treated with CX-4945. Biological processes significantly represented in phosphoproteomic profile were identified using annotations from GO database. The *p*-value of modified Fisher Exact Test from DAVID is placed in square brackets.

Recently, another phosphoproteomic study in non-small cell lung cancer (NSCLC) cell line NCI-H125 using the clinical-grade synthetic peptide CIGB-300, found mRNA processing and ribosome biogenesis as biological processes modulated after CK2 inhibition [26]. Protein folding, cytoskeleton organization, microtubule formation and protein ubiquitination were also significantly modulated after treatment with CIGB-300 [26]. According with both studies, CK2 inhibition by CX-4945 or CIGB-300 modulates a common set of biological processes but also each drug exerts its own mechanism of action by modulating a unique array of phosphoproteins. Since this effect could be a consequence of the different neoplastic backgrounds explored in each study (AML and NSCLC), a phosphoproteomics study of AML cells treated with CIGB-300 is currently underway to validate our hypothesis.

Noteworthy, proteins involved in cellular response to DNA damage appeared differentially phosphorylated in HL-60 cells treated with CX-4945 (**Figure 3**). Accordingly, CK2-mediated phosphorylation has been verified to regulate proteins with critical role in DNA damage response and DNA repair pathways [52]. In fact, phosphoproteomic analysis of cells treated with radiomimetic compound or ionizing radiation to induce DNA double-stranded breaks showed a dynamic response for a significant number of CK2 phosphorylation motifs [53, 54]. Furthermore, combination of CK2 inhibitors with DNA-targeted drugs evidenced a synergistic interaction in cancer models, owing to the suppression of DNA repair response triggered by such chemotherapeutic agents [55, 56]. Interestingly, a number of modulated phosphorylation sites in AML cells belong to proteins implicated in regulation of viral process (**Figure 3**). The relevance of CK2 in viral infections has been well documented, and a number of viral and cellular proteins essential for virus replicative cycle and pathogenesis are listed as *bona fide* CK2 substrates [57].

On the whole, CK2 inhibition with CX-4945 impacted on a broader set of biological processes in OCI-AML3, which is in agreement with the higher number of differentially modulated phosphopeptides in this cell line (**Figure 2A, Figure 3**). However, as pointed above such divergence does not impinge on the antiproliferative effect exerted by CX-4945.

### 2.3. Sequence Analysis of Phosphopeptides Identified in AML Cells

Protein kinases recognize structural and sequence motif, which in conjunction with other factors like subcellular co-localization or protein complex formation, determine their specificity [58]. Particularly, CK2 phosphorylation is specified by multiple acidic residues located mostly downstream from the phosphoacceptor amino acid, the one at position n + 3 playing the most crucial function. Besides, proline residue at position n + 1 acts as a negative determinant for protein kinase CK2 phosphorylation [3, 5].

In our study, approximately 21% of the phosphopeptides identified in HL-60 and OCI-AML3 fulfill the CK2 consensus sequence (**Figure 4A, Table S3**). This proportion of putative CK2 substrates is in accordance with previous phosphoproteomic analysis [24, 59]. In HL-60 the majority of phosphopeptides (83.3%) containing the CK2 consensus sequence were unaffected by CX-4945 treatment. Moreover, 107 phosphopeptides (16.7%) containing the CK2 consensus sequence were significantly modulated in HL-60 treated cells, of which 14.4% had a decreased and 2.3% had an increased phosphorylation respect to non-treated cells (**Figure 4A**). In contrast to HL-60 cells, the majority of phosphopeptides (53.9%) containing the CK2 consensus sequence had a decreased phosphorylation in OCI-AML3 cells treated with CX-4945, whereas 45.8% were unaffected and 0.3% had an increased phosphorylation (**Figure 4A**). This result reinforces the differential impact of CX-4945 over the CK2-dependent signaling, which was evidenced above by the higher number of total phosphopeptides that had a decreased phosphorylation in OCI-AML3 treated cells (1310 out of 2976) (**Figure 2A**).

**Figure 4.**
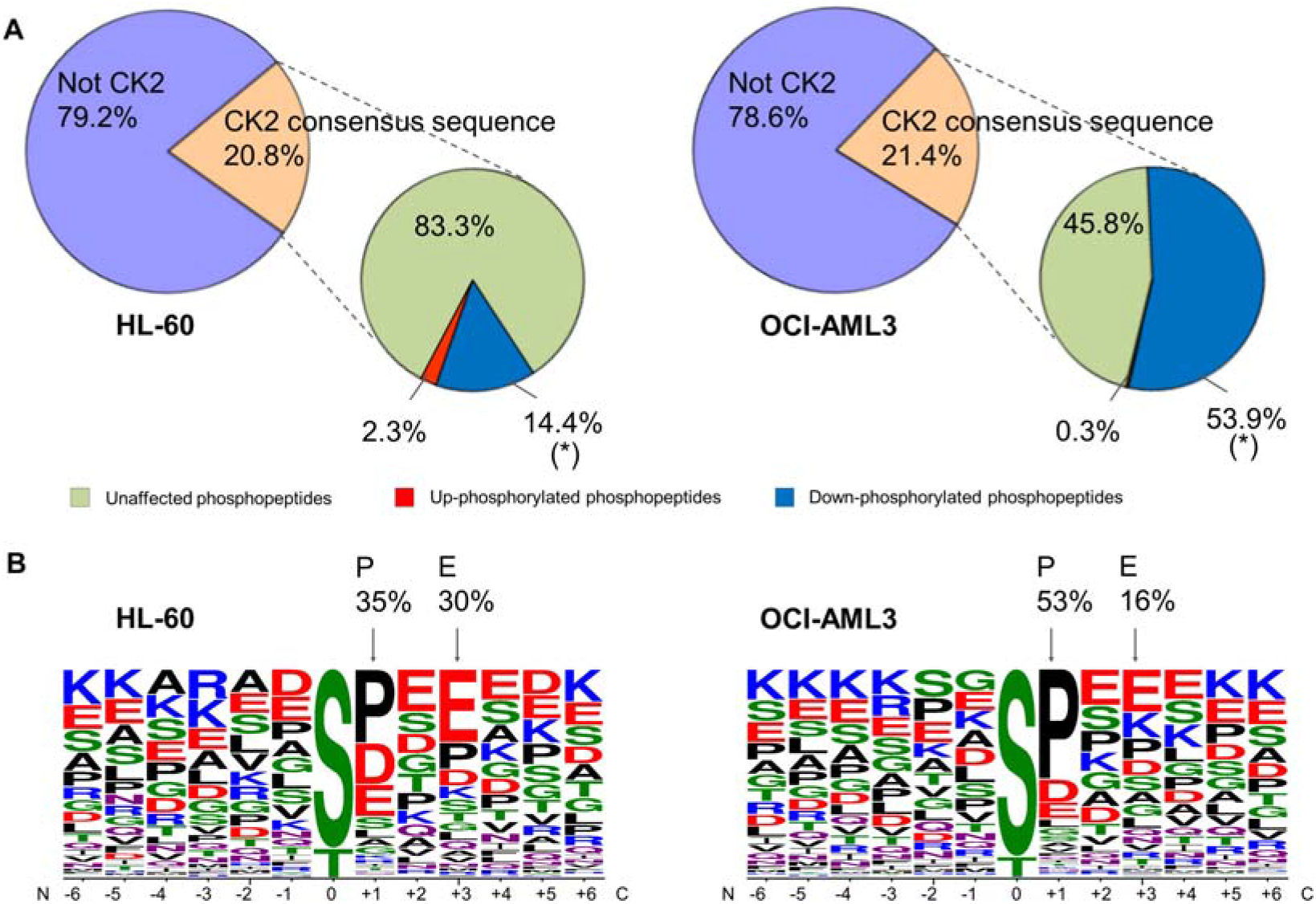
**S**equence analysis of phosphopeptides identified in AML cells treated with the CK2 inhibitor CX-4945: (**a**) Pie charts show the percent of phosphopeptides identified in HL-60 and OCI-AML3 cells that either, contains or not the CK2 consensus sequence. For the former category, the percentage of phosphopeptides that are significantly increased or decreased, or that do not show significant changes in their phosphorylation levels are reported in lateral pie charts; (**b**) Sequence logos corresponding to phosphopeptides significantly down-phosphorylated in AML cells treated with CX-4945. Logos were generated using WebLogo tool and MaxQuant amino acid sequence window as input [38]. (*) Phosphopeptides with decreased phosphorylation due to the reduction of protein abundance were not considered as differentially regulated.

CK2 substrates have different rates of phosphorylation turnover, some of them are promptly reduced after 6 h of treatment with CX-4945 but others are more resistant to dephosphorylation, since requires much longer treatment times (up to 24 h) and higher concentrations of the inhibitor [24]. We think that the foregoing could explain the proportion of putative CK2 phosphopeptides that resulted unaffected after 8 h of treatment with CX-4945 in AML cells. Even more, in C2C12 cells devoid of CK2 catalytic activity (CK2α/α′^(|/|)^) was demonstrated that not all the phosphopeptides conforming the CK2 consensus sequence have reduced phosphorylation levels, suggesting that other kinase(s) could fulfill the phosphorylation of these sites in the absence of CK2 [27].CK2 consensus is a quite distinctive motif where phosphoacceptor amino acid is surrounded by acidic residues [5]. As demonstrated by sequence logo analysis, the positions up- and down-stream of phosphorylated sites in peptides that significantly decreased after treatment with CX-4945 are predominantly occupied by acidic residues (**Figure 4B**). Furthermore, 30% and 16% of the phosphopeptides down-regulated by CX-4945 had a glutamic acid at position n + 3 in HL-60 and OCI-AML3 cells, respectively (**Figure 4B**). Basic residues are less represented or practically absent at positions spanning between n + 1 to n + 4. All these features are consistent with the previously reported linear motif preference of CK2.

Notably, phosphopeptides containing the S/T-P motif were also down-phosphorylated in AML cells after CK2 inhibition with CX-4945 (**Figure 4B**). In fact, 35% and 53% of the significantly down-phosphorylated peptides had a proline at position n + 1 in HL-60 and OCI-AML3 cells, respectively (**Figure 4B**). This motif is targeted by the large and heterogeneous category of proline-directed kinases and has been previously reported that such motif is incompatible with direct phosphorylation by CK2 [60]. Thus, the down-regulation of phosphopeptides containing S/T-P motif could be interpreted as off-target effect of CX-4945 or just an indirect result of CK2 inhibition, i.e. perturbations of other kinases involved in signaling networks where CK2 is also implicated. Considering that this effect has been associated not only to CX-4945, but also to others CK2 inhibitors [24-26], we reasoned that decrease in phosphorylation such phosphopeptides is just a consequence of signaling propagation following CK2 inhibition.

### 2.4. Network Analysis of Kinases Associated with AML Phosphoproteomic Profiles

To identify kinases responsible for the phosphoproteomic profile modulated in HL-60 and OCI-AML3 cells, an enzyme-substrate network was constructed using iPTMnet and KEA2 bioinformatic resources [39, 42]. A total of 37 differentially modulated phosphopeptides in HL-60 cells (|FC| ≥ 1.5, *p*-value < 0.05) were attributed to 31 kinases including CK2 with the higher number (10 phosphopeptides) (**Figure 5, Figure S2, Table S4**). A broader picture was observed in OCI-AML3 phosphoproteome, in which 207 differentially modulated phosphopeptides were associated to 73 kinases. As expected, CK2 enzyme was again among the most represented kinases with 29 phosphopeptides (**Figure 5, Figure S2, Table S4**). Kinases significantly associated with the phosphoproteomic profile were also identified using KEA2 bioinformatic tool [42]. In addition to CK2, members of the CDKs and MAPKs families like CDK1, CDK2, MAPK9 and MAPK14 were also significantly associated with the OCI-AML3 phosphoproteome (**Figure S2**). These results are in accordance with sequence logo analysis, which indicates that CK2 and proline-directed kinases motifs are the most frequent among the phosphopeptides down-regulated after CK2 inhibition in AML cells.

**Figure 5.**
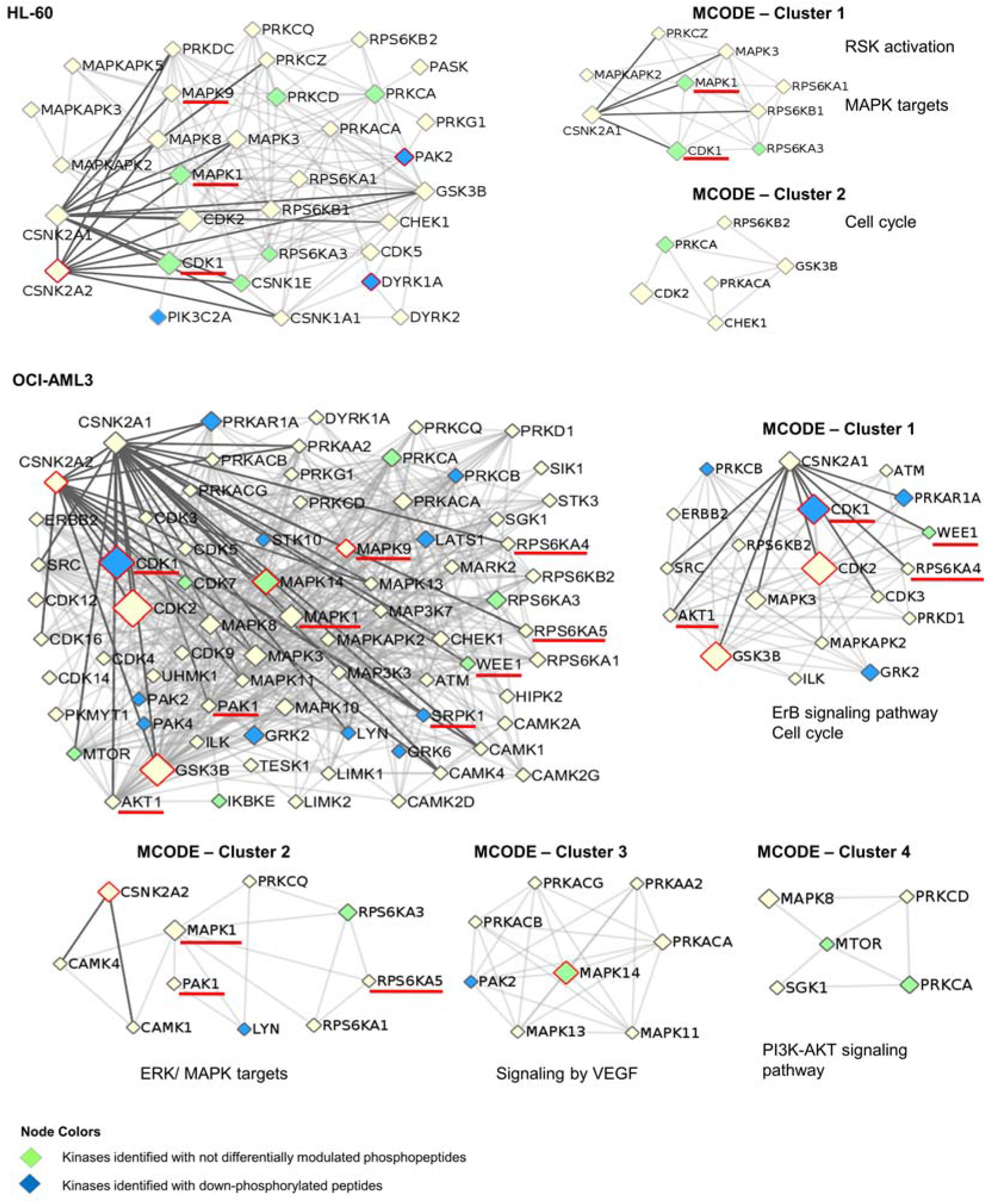
Protein-protein interaction network of kinases associated to phosphoproteomic profiles differentially modulated by CX-4945 in HL-60 and OCI-AML3 cells. Protein clusters were identified with MCODE algorithm and the related biological processes and signaling pathways are indicated. For each protein kinase the node size is proportional to the number of target phosphopeptides that appeared differentially phosphorylated in response to CK2 inhibition. Kinases that are significantly associated with the phosphoproteomic profiles, according to KEA2 results, are highlighted with a red border. In addition, kinases indicated with a red line are bona fide CK2 substrates, whereas green and blue nodes correspond to those kinases that were identified in our analysis with either not differentially modulated phosphopeptides and down-phosphorylated peptides, respectively.

An interaction network of protein kinases associated with the phosphoproteomic profile modulated in HL-60 and OCI-AML3 cells was represented using the Metascape bioinformatic software (**Figure 5**) [43]. The kinome network also includes those kinases that were identified in AML cells after CK2 inhibition, with either not differentially modulated phosphopeptides (green nodes) or down-phosphorylated peptides (blue nodes). For instance, the tyrosine-phosphorylated and regulated protein kinase DYRK1A is known to promote cell proliferation and survival [61]. DYRK1A is auto-phosphorylated in S529, modification that enhances 14-3-3-β protein binding and consequently increases the kinase catalytic activity [62]. DYRK1A S529 was found down-phosphorylated in our study, suggesting an inhibition of this kinase in HL-60 cells. In fact, the S369 of Cyclin-L2, a known DYRK1A substrate which is involved in RNA processing of apoptosis-related factors [63], was also found down-phosphorylated in HL-60 cells (**Figure S2**).

CK2 has direct interactions with 13 and 27 kinases related to the phosphoproteomic profile identified in HL-60 and OCI-AML3 cells, respectively (**Figure 5**). Such kinases include nine *bona fide* CK2 substrates, three of them (MAPK1, MAPK9 and CDK1) related to both phosphoproteomics profiles (**Figure 5**). Although none of the CK2 phosphosites belonging to these kinases were identified in the present study, the results suggest a signal propagation downstream of these proteins. For instance, CK2 phosphorylates mitogen-activated protein kinase 1 (MAPK1) at S246 and S248, such event promotes MAPK1 nuclear translocation and phosphorylation of target transcription factors [64]. A total of 19 phosphopeptides which are substrates of MAPK1 were identified down-phosphorylated in OCI-AML3 after CK2 inhibition (**Figure S2**). Besides, CK2 phosphorylates cyclin-dependent kinase 1 (CDK1) at S39 and regulates cell cycle [65]. Accordingly, the enzyme-substrate network evidenced an inactivation downstream of CDK1 since at least, 43 phosphosites modulated by CDK1 were down-phosphorylated in OCI-AML3 cells. Such phosphopeptides belong to proteins related to chromatin remodeling, mitotic spindle assembly, and DNA repair (**Figure S2**).

Highly connected regions in the kinome networks associated to HL-60 and OCI-AML3 phosphoproteomic profiles were identified using MCODE algorithm [44]. Clusters representing cell cycle and MAPK targets appeared as a common denominator in kinome networks from both AML cell lines (**Figure 5**). In contrast, signaling pathways mediated by VEGF and PI3K/AKT only appeared in OCI-AML3 kinome network (**Figure 5**). Protein kinase CK2 it is known that up-regulates PI3K/AKT pathway, in part by phosphorylating and activating AKT1 [66]. To note, PI3K/AKT pathway is constitutively active and sustain viability of primary acute lymphoblastic leukemia cells (ALL), signaling alteration that results from CK2 overexpression and hyperactivation [67]. AML and ALL are hematological diseases with several features in common, and previous studies have showed that the antineoplastic effect of CX-4945 in both malignancies is mediated by attenuation of the PI3K/AKT pathway [51, 68-70]. Accordingly, we found a number of AKT1 substrates down-phosphorylated in OCI-AML3 cells after CK2 inhibition with CX-4945, whereas in HL-60 cells the PI3K/AKT pathway did not appeared significantly represented in our analysis, explaining perhaps the refractoriness to CX-4945-induced apoptosis displayed by this cell line.

Importantly, previous phosphoproteomic results from primary AML cells have indicated that at the diagnosis time, patients that relapse after chemotherapy had a higher CK2, MAPK and CDK activity in comparison with patients which have free-relapse evolution [22]. However, the high CK2 activity at diagnosis of relapsed patients was no longer observed in chemoresistant cells [23]. Aasebø *et al*. pointed out that the proteome and phosphoproteome profiles changed considerably from the first diagnosis to the first relapse, therefore CK2 could be important in inducing treatment-resistant clones but dispensable for the survival of clones that already have become resistant to therapy [23]. Remarkably, in our study substrates of CK2, MAPKs and CDKs were found down-phosphorylated after CX-4945 treatment of AML cell lines, being MAPKs and CDKs signaling modulation probably a down-stream consequence of CK2 inhibition (**Figure 5, Table S4**).

### 2.5. Identification of CK2 Substrates Modulated by CX-4945 in AML Cells

Besides the *bona fide* CK2 substrates annotated in iPTMnet and KEA databases [39, 42], additional candidate CK2 substrates in AML cells were searched. According to the presence of the CK2 consensus sequence, 39% and 26% of all differentially modulated phosphopeptides on HL-60 and OCI-AML3 respectively, could be putative CK2 substrates responsive to CX-4945. However, phosphosites recognized by other protein kinases like Ser/Thr-protein kinase Chk1 or cAMP dependent protein kinase catalytic subunit alpha (PKACA) could contain an acidic amino acid at position n + 3 (**Figure S3**). Indeed, we observed that arginine is frequent at position n – 3 from the phosphorylated residue (**Figure 4**), a motif that is recognized by basophilic kinases [59]. Therefore, we search for additional evidences in support phosphoproteins containing the CK2 consensus sequence as candidate CK2 substrates.

First, differentially phosphorylated proteins identified in AML cells were searched as candidate CK2 substrates using NetworKIN database [45]. Such database includes enzyme-substrate interactions predicted not only based on the consensus sequence recognized by the enzyme, but also using a protein association network to model the context of substrates and kinases, which improves the prediction accuracy [45]. Second, the phosphoproteomic profile differentially modulated in AML cells after CK2 inhibition was compared with a dataset of high confidence CK2 substrates reported by Bian *et al*. [46]. These authors identified *in vitro* CK2 substrates by combining kinase reaction on immobilized proteomes with quantitative phosphoproteomics, and to reduce false positive results compared *in vitro* phosphosites with *in vivo* phosphorylation sites reported in databases [46]. Lastly, the differentially modulated phosphoproteins that interact with CK2 were searched using Metascape, which performed interactome analysis based on integrative protein-protein interactions databases like InWeb_IM and OmniPath [43].

Taking into account the four levels of predictions (CK2 consensus sequence, NetworKIN prediction, CK2 substrates predicted by Bian *et al*. [46] and interaction with CK2) we identified a total of 117 and 359 candidate CK2 substrates differentially modulated after CK2 inhibition in HL-60 and OCI-AML3 cells, respectively (**Table S5**). This dataset was filtered out to find those substrates that had the concomitant occurrence of two or more criteria associated to CK2 phosphorylation. Applying this workflow, in HL-60 cells 64 phosphosites on 53 proteins were identified as the most reliable CK2 substrates modulated after treatment with CX-4945, whereas 168 phosphosites on 120 proteins were identified in OCI-AML3 cells (**Figure 6, Table S5**). The list includes those CK2 substrates previously confirmed as *bona fide* according to iPTMnet and KEA databases [39, 42].

**Figure 6.**
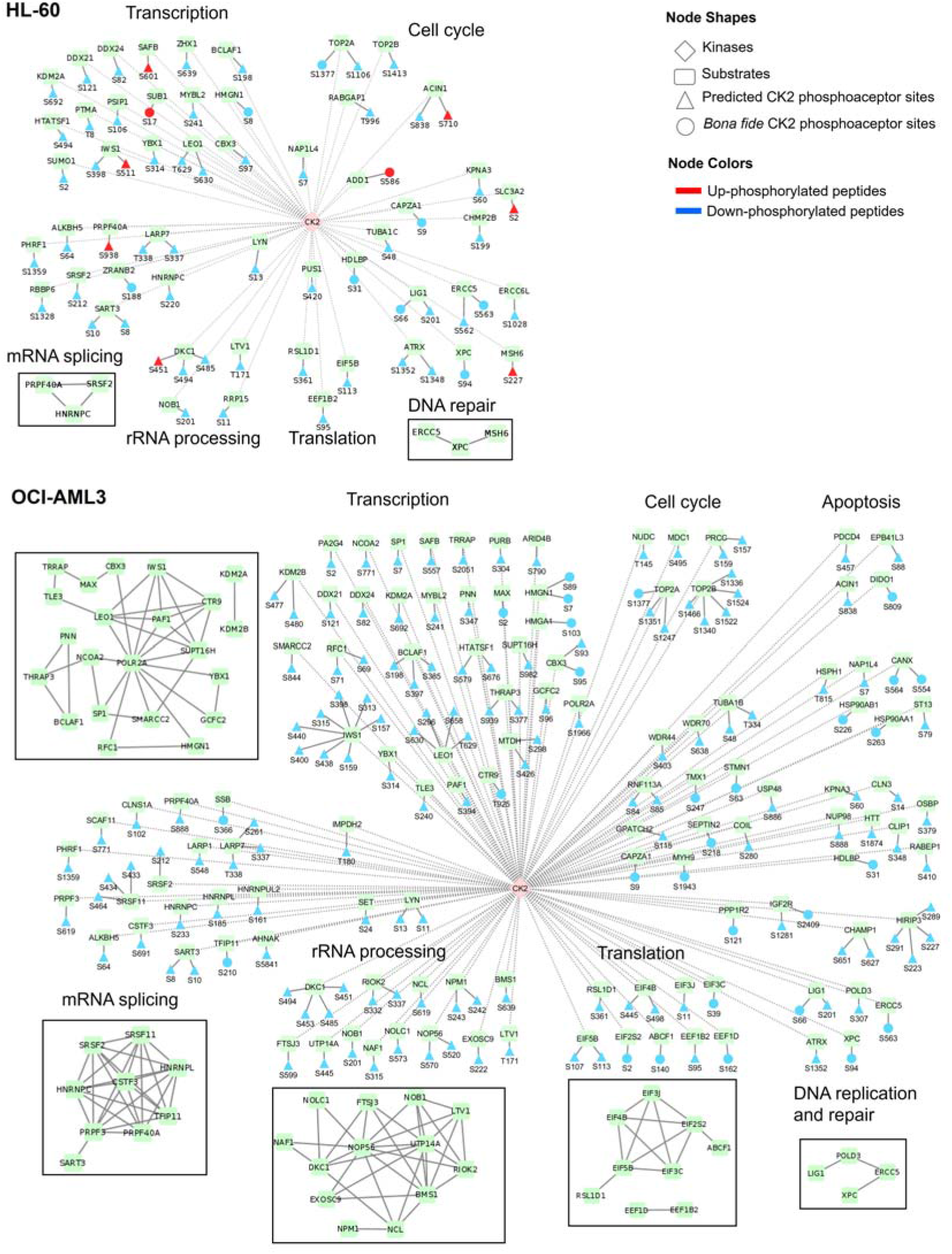
Network of CK2 substrates differentially modulated after CK2 inhibition with CX-4945 in AML cells. For each substrate, the phosphoacceptor sites (*bona fide* and predicted) for CK2-mediated phosphorylation and its modulation after incubation with CX-4945 are indicated. Phosphoproteins are grouped according to related biological processes annotated in GO database and squares representing protein-protein interactions networks retrieved from STRING database are shown.

Remarkably, for the 67% and 71% of the high confidence CK2 substrates modulated in HL-60 and OCI-AML3 cells, respectively, any related enzyme was annotated in iPTMnet database. Besides, to our knowledge the phosphosites S280 of coilin protein and T180 of inosine-5’-monophosphate dehydrogenase 2 (IMPDH2) are reported for the first time. Coilin protein is an integral component of Cajal bodies-subnuclear compartments, whereas IMPDH2 catalyzes the first and rate-limiting step for *de novo* guanine nucleotide biosynthesis pathway [71, 72]. Interestingly, both proteins regulate cell growth and have been related to malignant transformation [72, 73]. However, validation of coilin S280 and IMPDH2 T180 as phosphorylation sites targeted by CK2 and the biological roles of such post-translational modifications need further experimentation.

### 2.6. Functional Characterization of CK2 Substrates Identified in AML Cells

Phosphoproteins identified as candidate CK2 substrates are related to transcription, mRNA splicing, rRNA processing, translation, DNA repair and cell cycle in both AML cells lines (**Figure 6**). However, the number of potential CK2 substrates differentially modulated after CK2 inhibition is higher in OCI-AML3 cells than in HL-60 cells. As pointed before, this could explain the different sensitivity to CX-4945 cytotoxic effect of HL-60 cells in comparison to other AML cell lines [51]. In fact, we identified candidate CK2 substrates related to apoptosis only in the phosphoproteomic profile of OCI-AML3 cells (**Figure 6**). This subset includes three tumor suppressors: erythrocyte membrane protein band 4.1 like 3 (EPB41L3 S88), the programmed cell death 4 protein (PDCD4 S457) and the death inducer-obliterator 1 (DIDO1 S809). However, the effect of CK2-mediated phosphorylation for the function of these proteins remains to be determined.

CK2 inhibition in AML cells could impact the transcriptional machinery by modulating the phosphorylation of several candidate substrates. Such CK2 candidate substrates in OCI-AML3 phosphoproteomic profile are centered around the RNA polymerase II subunit A (POLR2A) according to protein-protein interactions gathered from STRING database (**Figure 6**) [47]. Three components of the PAF1 complex which interacts with RNA polymerase II during transcription were identified as candidate CK2 substrates: RNA polymerase II-associated factor 1 homolog (PAF1 S394), RNA polymerase-associated protein LEO1 (LEO1 S296, S630, S658 and T629) and RNA polymerase-associated protein CTR9 homolog (CTR9 T925). PAF1 complex is required for transcription of Hox and Wnt target genes [74]. Therefore, down-phosphorylation of these candidate substrates could modulate the Wnt signaling pathway. Supporting this hypothesis, previous studies highlights that CK2 is a positive regulator of Wnt signaling pathway and CK2 inhibition by CX-4945 has been associated with Wnt/β-catenin inhibition [75, 76].

Substrates related to transcription include *bona fide* CK2 targets such as the non-histone chromosomal protein HMG-14 (HMGN1) and the high mobility group protein HMG-I/HMG-Y (HMGA1) [77-79]. The phosphorylation level of both proteins (HMGN1 S7, S8, S89; HMGA1 S103) decreased after CK2 inhibition by CX-4945 (**Figure 6**). Importantly, AML patients that relapsed after chemotherapy have an increased phosphorylation level of HMGN1 S7 [22]. In general HMG proteins modulate chromatin and nucleosome structure, participate in transcription, replication, DNA repair, and extracellular HMGN1 has been described to function as an alarmin that contributes to the generation of innate and adaptative immune responses [80, 81]. The biological effect of CK2 phosphorylation of HMGN1 and HMGA1 is currently unknown, although, previous studies suggest that phosphorylation of HMGN1 could interfere with its nuclear localization [78].

The most densely down-phosphorylated protein among the candidate CK2 substrates is the protein IWS1 homolog (IWS1) which was identified with eight phosphopeptides in OCI-AML3 cells (**Figure 6**). This protein recruits a number of mRNA export factors and histone modifying enzymes to the RNA polymerase II elongation complex and modulates the production of mature mRNA transcripts [82, 83]. As illustrated by **Figure 6**, several candidate CK2 substrates related to mRNA splicing were down-phosphorylated after CK2 inhibition in AML cells, including members of the spliceosome complex. Among those proteins are heterogeneous nuclear ribonucleoproteins (HNRNPC, HNRNPL), serine and arginine rich splicing factors (SRSF2, SRSF11) and pre-mRNA processing factors (PRPF3 and PRPF40A) (**Figure 6**). In particular, CK2 phosphorylation of heterogeneous nuclear ribonucleoproteins C1/C2 (HNRNPC) it known that regulates its binding to mRNA [84, 85]. In agreement with our results, was previously demonstrated that CK2 inhibition by quinalizarin and CIGB-300 modulates a subset of CK2 substrates related to transcription, RNA processing and mRNA splicing [24, 26]. To note that at the time of diagnosis, phosphoproteins containing CK2 phosphoacceptor sites and related to RNA processing have an increased phosphorylation level in relapse AML patients when compared to those which have a relapse-free evolution [22]. Another phosphoproteomic study comparing pairing samples of AML patients at the time of diagnosis and first relapse found that also RNA-splicing and -binding proteins were up-phosphorylated at first relapse [23].

CK2 phosphorylation of proteins related to rRNA processing and translation has been well documented [3]. Among the proteins probably subject to CK2 regulation in AML cells are members of the nucleolar ribonucleoprotein complex (NAF1 S315; DKC1 S451, S453, S485, S494; NOP56 S520, S570) (**Figure 6**). According to information gathered from STRING database [47], such proteins interacts with phosphoproteins related to ribosome biogenesis (RIOK2 S332, S337; BMS1 S639; LTV1 T171) which were identified mainly in OCI-AML3 cells (**Figure 6**). The effect of CK2 regulation of these proteins remains to be elucidated. However, the results highlight the important role of CK2 in regulating protein biosynthesis to support the high proliferative rate of tumor cells. In line with this result, a cluster of eukaryotic translation initiation factors (EIF) was down-phosphorylated after CK2 inhibition (**Figure 6**). This cluster contains two members of the EIF3 complex: EIF3J S11 and EIF3C S39. EIF3J is a known CK2 substrate and its phosphorylation on S127 promotes assembly of EIF3 complex and activation of the translational initiation machinery [86]. Besides, CK2 phosphorylates EIF2β on S2, a phosphopeptide also identified in our study, and such modification stimulates EIF2β function in protein synthesis [87]. Down-phosphorylation of proteins related to the translational machinery after CK2 inhibition could add a beneficial impact at the clinical evolution of AML patients, since protein translation has been associated with increased relapse risk [22, 23].

Another function attributed to CK2 is the regulation of the cellular DNA damage response [52]. After CK2 inhibition in AML cells, the biological process of DNA repair appeared significantly represented in the phosphoproteomic profiles (**Figure 3**). A recent study demonstrated that proteins related to DNA repair have increased phosphorylation levels in relapse AML patients [22]. Among those phosphoproteins associated with such unfavorable chemotherapy outcome, we identified in our study that treatment of AML cells with CX-4945 down-phosphorylates TRIM28 S19, TP53BP1 S523/S525 and LIG1 S66, this latter a known CK2 substrate (**Table S1**) [88]. Besides, others known and putative CK2 substrates related to DNA repair were also found down-phosphorylated in our study, like the DNA damage recognition and repair protein (XPC S94) (**Figure 6**). In particular, CK2 phosphorylation of XPC at S94 promotes recruitment of ubiquitinated XPC to the chromatin which is important for nucleotide excision repair following ultraviolet induced DNA damage [89]. Previous studies demonstrated that CK2 inhibition by CX-4945 inactivates the function of other essential DNA repair proteins, supporting the synergistic interaction of this inhibitor with chemotherapeutic agents that induce DNA damage [55].

Worthy of note, we identified members of the heat shock protein 90 (HSP90) chaperone proteins differentially modulated in OCI-AML3 phosphoproteomic profile. CK2 mediated phosphorylation of HSP90 is required for its chaperone activity toward client kinases, some of them involved in human cancers [90, 91]. Phosphosites from HSP90-alpha (HSP90AA1 S263) and HSP90-beta (HSP90AB1 S226) were both down-phosphorylated after CK2 inhibition in OCI-AML3 cells (**Figure 6**). Thus, modulation of HSP90 by CX-4945 in OCI-AML3 cells could be in part responsible for the signal propagation downstream of CK2 inhibition and the pronounced effect over the kinome network in this cell line. In agreement with our findings, besides attenuation of PI3K/AKT pathway, disruption of unfolded protein response (UPR) have also been pointed as a mediator of CX-4945-induced apoptosis in ALL cell lines and primary lymphoblasts [69, 70]. Importantly, in such effect the reduction of chaperoning activity of HSP90 appears to play a critical role [69, 70]. Moreover, in multiple myeloma (MM) cells, another hematological malignancy having common features with AML, has been documented that CK2 inhibition causes apoptotic cell death through alterations of the UPR pathway [92].

In summary we found that the phosphoproteomic profiles modulated after CK2 inhibition with CX-4945 in AML cell lines, contain protein mediators of signaling pathways and biological processes previously described in primary AML cells (**Figure 7**) [22, 23, 51, 68]. Therefore, our findings, in conjunction with Quotti Tubi *et al*. results and AML patients phosphoproteomic data from Aasebø *et al*., support the rationale of protein kinase CK2 pharmacologic inhibition for AML targeted therapy, an approach that could significantly improve the outcome in AML therapeutics.

**Figure 7.**
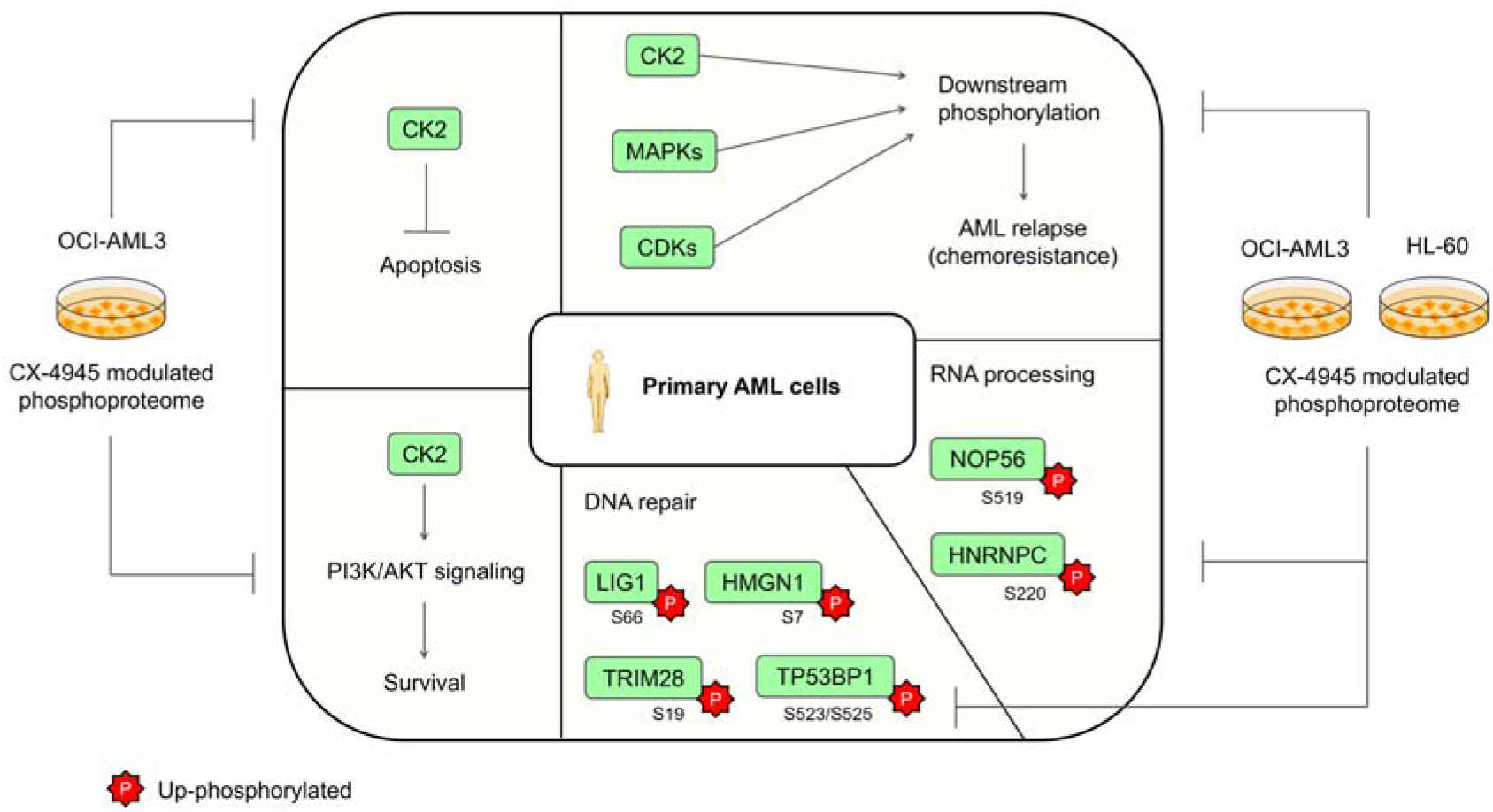
**S**ignaling pathways and biological processes deregulated in primary AML cells and modulated by the CK2 inhibitor CX-4945 in AML cell lines. Phosphoproteins up-regulated in primary AML cells and down-phosphorylated in CX-4945-treated AML cells are indicated.

## 5. Conclusions

Our study provides the first quantitative phosphoproteomic analysis exploring the molecular impact of the ATP-competitive CK2 inhibitor CX-4945 in human cell lines representing two differentiation stages and major AML subtypes. Here, we identified a total of 273 and 1310 unique phosphopeptides as significantly modulated in HL-60 and OCI-AML3 cells, respectively. Modulated phosphopeptides are mainly related to mRNA processing and splicing, response to DNA damage stimulus, protein sumoylation and regulation of viral processes. In addition, the network analysis illustrated how the relationship of CK2 with other kinases could orchestrate the perturbation of AML cells phosphoproteome. In this complex cellular response, phosphorylation mediated by other kinases besides CK2 could be interpreted as a consequence of signal propagation downstream of CK2 inhibition, rather than off-targets effects. Additionally, using database mining and prediction tools, in HL-60 cells we identified 64 phosphosites on 53 proteins as high confidence CK2 substrates responsive to CX-4945, whereas 168 phosphosites on 120 proteins were identified in OCI-AML3 cells. Such substrates not only explain the variety of cellular effects exerted by CX-4945, but also reinforce the instrumental role of protein kinase CK2 in AML biology. Finally, our results, in conjunction with previous findings in primary AML cells, support the suitability of using CK2 inhibitors for AML targeted therapy.

## Author Contributions

Conceptualization, S.E.P., Y.P., Y.R. and L.J.G.; methodology, J.R.W. and K.Z.; software, O.G.; formal analysis, M.R., A.R. and A.C.R.; investigation, V.B. and G.V.P.; data curation, V.B.; writing-original draft preparation, M.R. and A.R.; writing-review and editing, A.C.R. and G.V.P.; supervision, S.E.P. and Y.P.; project administration, S.E.P.; funding acquisition, J.R.W. All authors have read and agreed to the version of the manuscript.

## Acknowledgments

We thank Igor Parón for the support in mass spectrometry analysis.

## Figures and Supplementary Materials

**Figure S1.**
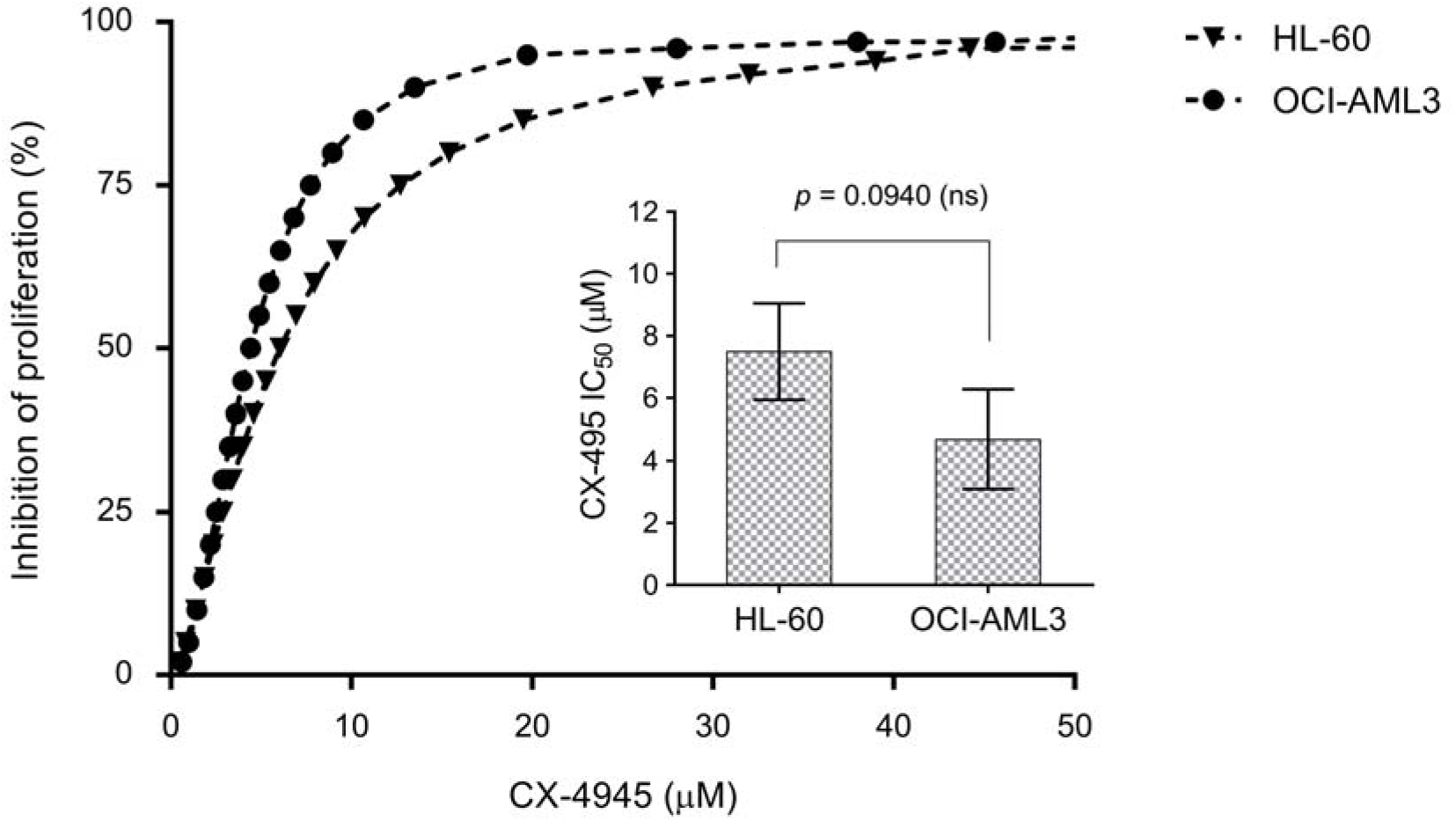
Antiproliferative effect of CK2 inhibitor CX-4945 in AML cell lines. Proliferation of HL-60 and OCI-AML3 cells was assessed using AlamarBlue assay. Dose-response curves are representative of three independent experiments and IC_50_ values are shown as mean ± SD, n = 3. ns, not significant.

**Figure S2.**
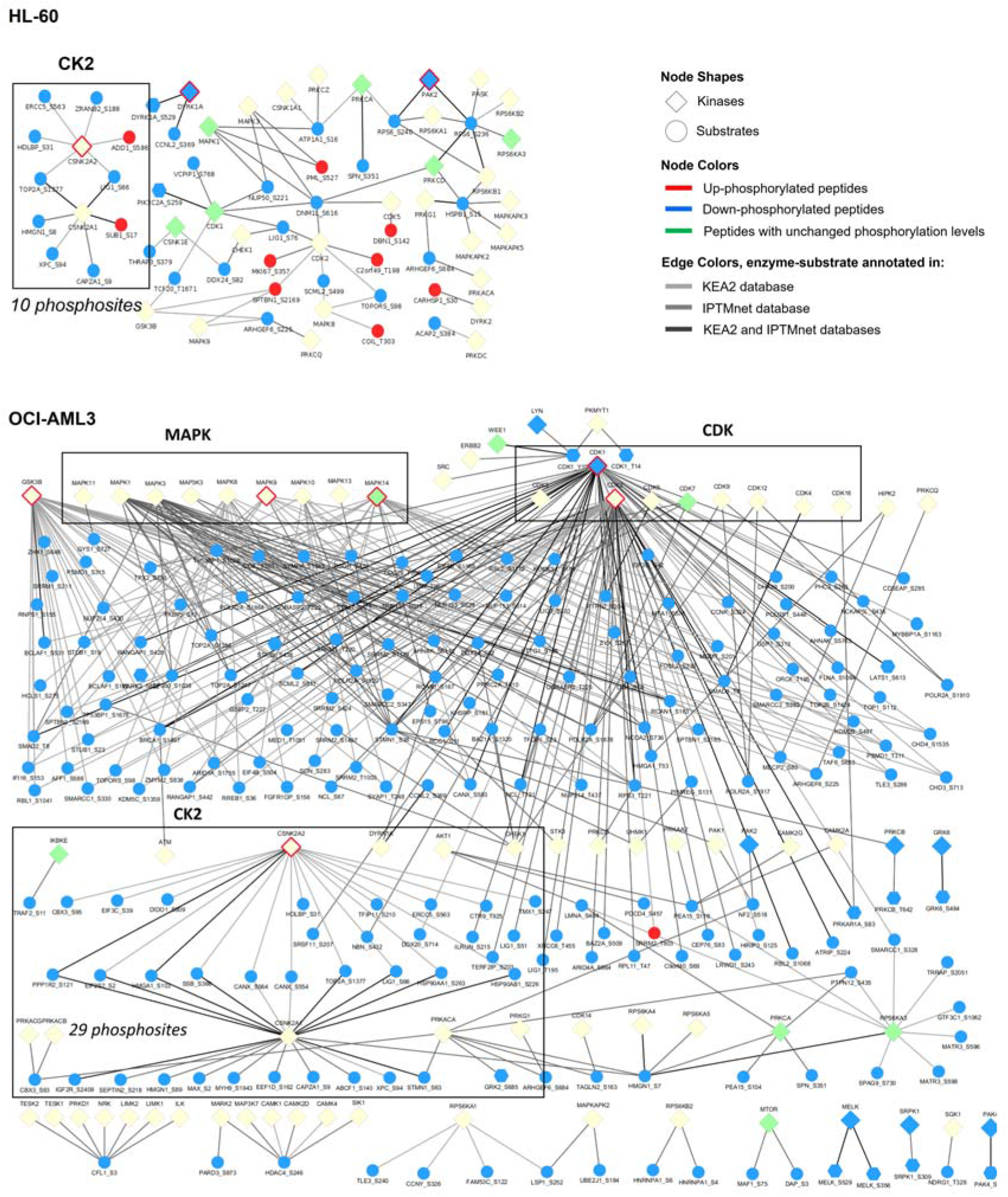
Enzyme-substrate network of differentially modulated phosphopeptides identified in AML cells using annotations from iPTMnet and KEA2. In squares are indicated the MAPK and CDK families as well as the CK2 substrates differentially modulated in HL-60 and OCI-AML3 cells after CK2 inhibition by CX4945. Kinases that are significantly associated with the phosphoproteomic profiles, according to KEA2 results, are highlighted with a red border. Network includes all kinases associated to AML cells modulated phosphoproteome, however green and blue nodes correspond to those kinases that were identified in our analysis with either not differentially modulated phosphopeptides and down-phosphorylated peptides, respectively.

**Figure S3.**
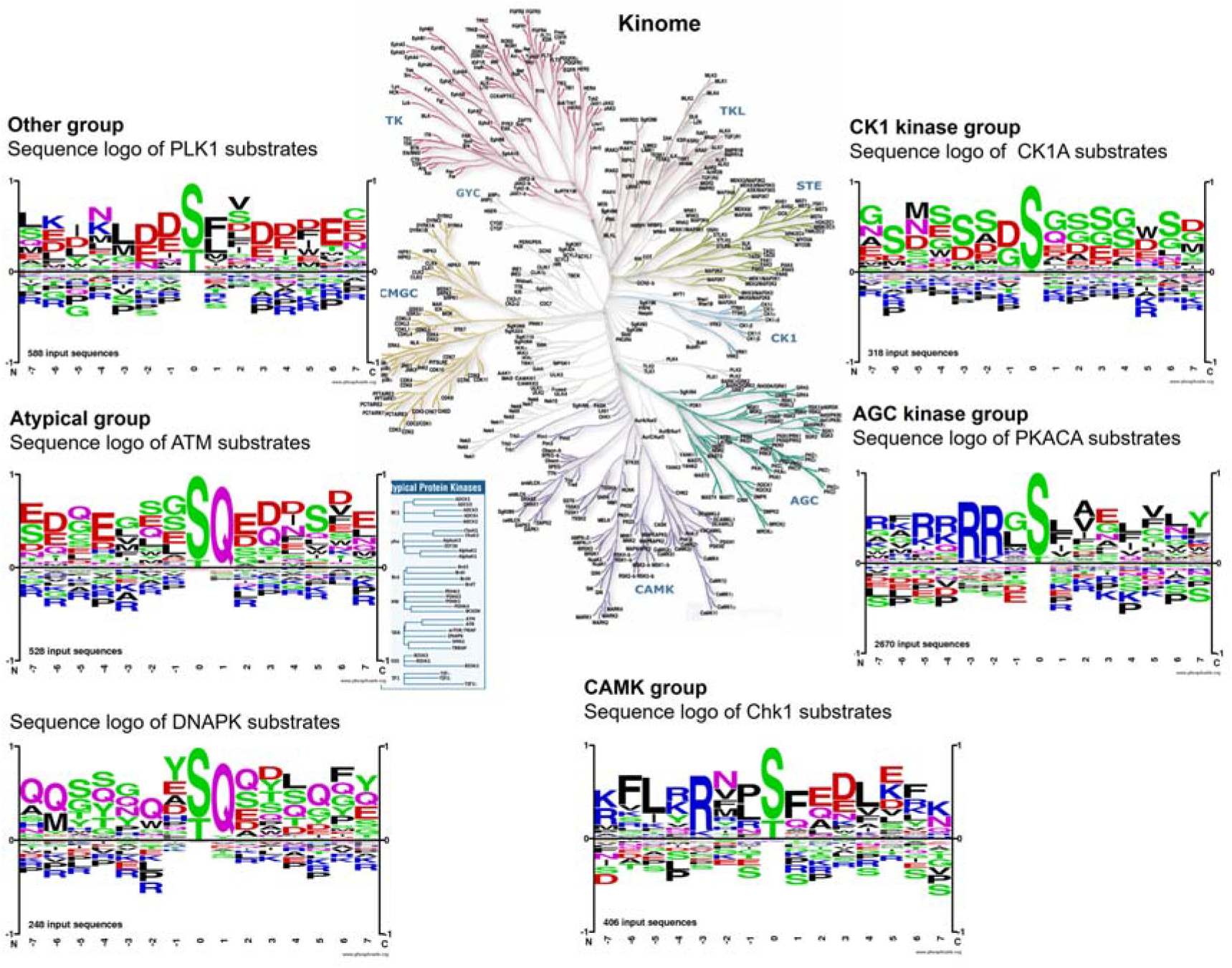
Sequence logos of phosphopeptides targeted by protein kinases representing five kinase groups (CAMK, Atypical, CK1, AGC and other) in the human kinome. The analysis was performed using the Motif Analysis tool of PhosphositePlus (PSP) (http://www.phosphosite.org/). All the substrates annotated in PSP database for each kinase were used as input sequences.

**Table S1**. Phosphoproteomic profile of AML cells treated with the CK2 inhibitor CX-4945.

**Table S2**. Proteins differentially modulated in AML cells treated with the CK2 inhibitor CX-4945.

**Table S3**. Phosphopeptides that fulfill the CK2 consensus sequence in AML phosphoproteomic profiles.

**Table S4**. Data mining of kinases associated to differentially phosphorylated peptides in AML phosphoproteomic profiles.

**Table S5**. Candidate CK2 substrates differentially modulated in AML cells treated with CX-4945.

